# Direct cleavage of human NLRP1 by enteroviral 3C protease triggers inflammasome activation in airway epithelium

**DOI:** 10.1101/2020.10.14.325076

**Authors:** Kim S. Robinson, Tan Kai Sen, Daniel Eng Thiam Teo, Ong Hsiao Hui, Bijin Au, Chrissie Lim, Lew Tian Sheng, Justin Chu Jang Hann, Vincent Tak Kwong Chow, Wang De Yun, Franklin L. Zhong, Bruno Reversade

## Abstract

Viruses pose a constant threat to human health. As a result our innate immune system has evolved multiple strategies to detect the presence of intracellular viral ‘pathogen-associated molecular patterns’ (PAMPs) (*1*). The full repertoire of human immune sensors and their PAMP ligands are not completely understood. Here we report that human NLRP1 senses and is activated by 3C proteases (3Cpros) of enteroviruses. Mechanistically, 3Cpros cleave human NLRP1 at a single site immediately after its primate-specific PYRIN domain, leading to oligomerization of its C-terminal fragment. Expression of 3Cpros in primary human cells cause NLRP1-dependent ASC oligomerization, pyroptotic cell death and IL-1 secretion. Consistent with our observation that NLRP1 is the predominant endogenous inflammasome sensor in human airway epithelium, we find that its genetic deletion, or that of ASC, abrogates IL-18 secretion from rhinovirus (HRV)-infected primary human bronchial epithelial cells. Our findings identify the first cognate PAMP ligand for human NLRP1 and assign a new function for the NLRP1 inflammasome in human antiviral immunity and airway inflammation. These results challenge the widely held notion that viral proteases largely serve to disable host immune sensing, and suggest that the human NLRP1 inflammasome may be a therapeutic target to treat inflammatory airway diseases including asthma.

**One Sentence Summary:** Human NLRP1 is activated by enteroviral 3C proteases

## Main Text

The human innate immune system employs a multitude of germline-encoded sensor proteins to detect microbial infections and kickstart the cell-intrinsic immune response (*2, 3*). Nod-like receptor (NLR) proteins are a family of innate immune sensors that can detect PAMPs in the cytosol (*3*–*5*). Upon activation, NLR proteins nucleate the assembly of distinct inflammasome complexes, leading to pyroptotic cell death and secretion of pro-inflammatory cytokines, such as IL-1B and IL-18 (*6*). Among all the NLR sensors found in humans, NLRP1 remains one of the least characterized. Germline activating mutations in *NLRP1*cause a Mendelian syndrome characterized by multiple self-healing keratoacanthomas and generalized hyperkeratosis (*7*–*9*), while common *NLRP1*single nucleotide polymorphisms (SNPs) are genetic risk factors for inflammatory and auto-immune diseases, such as asthma and vitiligo (*10, 11*). Human NLRP1 differs from rodent homologues in terms of domain organization, ligand specificity and tissue distribution (*12*). To date, no clinically-relevant pathogens or conserved PAMPs have been shown to specifically activate the human NLRP1 inflammasome.

Certain rodent Nlrp1 homologs, such as murine Nlrp1b can be activated by the *Bacillus anthracis*lethal factor toxin (LF), which directly cleaves Nlrp1b close to its N-terminus (*13*–*16*) (Fig.1A). The consensus LF cleavage site is conspicuously absent in human NLRP1, which contains a N-terminal PYRIN domain not found in rodents (Fig. 1A). As such, LF cannot cleave and activate human NLRP1. We hypothesized that the recently described mechanism of ‘activation by cleavage and functional degradation’ for murine Nlrp1b might still hold true for human NLRP1 and, if so, the responsible PAMP ligand might be derived from a human-specific pathogen (*3, 15, 16*). Several recent publications drew our attention to the human rhinovirus (HRV) (*17, 18*), the causative agent for the common cold. HRV is a member of the enterovirus genus (family: *Picornaviridae*), a family of ssRNA viruses that cause a range of human diseases, such as hand-foot-and-mouth disease, peri/myocarditis and poliomyelitis (*19*). HRV infection of primary human bronchial epithelial cells has been shown to induce caspase-1 activation and IL-1 secretion, although the upstream viral sensing mechanism has not been documented (*18, 19*). Notably, all picornaviruses, including HRVs, encode two well-defined proteases termed 2A and 3C, which are essential biogenesis factors required to cleave viral genomic precursor proteins into individual viral components (*20*). Both proteases also cleave a range of host cellular proteins (*21*–*24*) during infection of human cells.

**Fig. 1.**
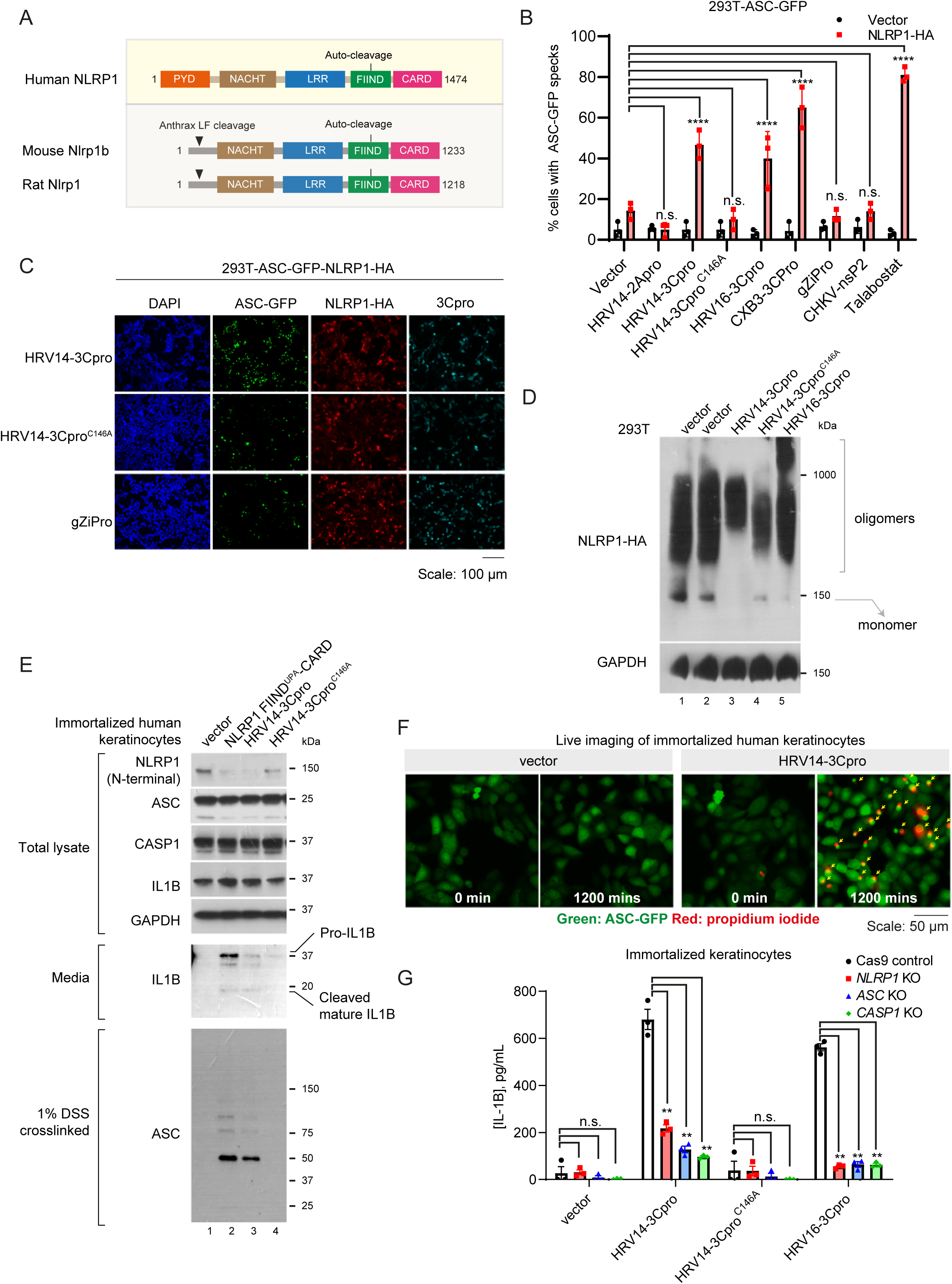
3C proteases activate the human NLRP1 inflammasome. A. Domain structures of human NLRP1 and rodent Nlrp1 homologues. B. Percentage of 293T-ASC-GFP cells with ASC-GFP specks after over-expression of vector control or NLRP1 with various viral proteases. Cells were fixed 24 hours after co-transfection of the indicated plasmids and >100 cells were scored for ASC-GFP speck formation. n=3 biological replicates. ****: p<0.0001 (two-way ANOVA). N.s: p>0.5 (two-way ANOVA). C. Representative images of ASC-GFP speck formation in NLRP1-expressing 293T-ASC-GFP cells transfected with the indicated viral proteases. D. NLRP1 oligomerization assayed by Blue-Native PAGE. 293T cells were transfected with NLRP1-HA and 3Cpros. Cells were lysed 48 hours after transfection. 20 μg of total lysate was used for Blue-Native PAGE followed by Western blot. E. HRV14-3Cpro induces mature IL-1B secretion and ASC oligomerization in immortalized human keratinocytes. N/TERT keratinocytes and conditioned media were harvested 48 hours after transfection. Conditioned media were concentrated 10 times before SDS-PAGE and IL-1B Western blotting. Endogenous ASC oligomers were extracted by 1% SDS after covalent crosslinking of the detergent (1% NP40)-insoluble pellets with 1% DSS. F. Live-cell imaging of ASC-GFP speck formation and lytic cell death in 3Cpro-transfected immortalized N/TERT keratinocytes. Yellow arrows, ASC-GFP specks. G. CRISPR/Cas9-deletion of *NLRP1, ASC*and *CASP1*abrogates 3Cpro-induced IL-1B secretion in immortalized human keratinocytes. Cas9-control, *NLRP1, ASC*and *CASP1*knockout keratinocytes were transfected with the indicated 3Cpro plasmids in parallel. Conditioned media were harvested 48 hours post transfection and analyzed by ELISA. **: p<0.005 (Two-way ANOVA), n=3 biological replicates.

To test whether these two proteases were sufficient to activate human NLRP1, we co-expressed NLRP1, HRV14-3C and HRV14-2A proteases in a 293T-ASC-GFP reporter cell line. While the HRV14 2A protease did not cause an increase in ASC-GFP speck formation relative to vector control, HRV14-3Cpro robustly induced ASC-GFP speck formation in the presence of NLRP1 (Fig. 1B-C). 3Cpros from another HRV strain (HRV16) and other enteroviruses, such as coxsackie A16 and B3, were all capable of inducing ASC-GFP speck formation to varying degrees in an NLRP1-dependent manner (Fig. 1B,fig. S1A-B). In comparison, neither the Zika (a flavivirus) NS2B-3 protease nor the Chikungunya (an alphavirus) nsP2 protease was able to induce ASC-GFP speck formation (Fig. 1B-C, fig. S1B). Furthermore, the ability of HRV14-3Cpro to activate human NLRP1 was entirely dependant on its enzymatic activity, since mutating its catalytic cysteine residue (p. C146A) completely abrogated ASC-GFP speck formation (Fig. 1B-C). In addition, transient co-expression of HRV14-, HRV16- and CXB3-3Cpros with human NLRP1 significantly increased the amount of high-molecular weight NLRP1 oligomers at the expense of monomeric NLRP1 (Fig. 1D,lanes 3, 5 and 6 vs. lane 1 and 2), suggesting that 3Cpros can directly activate NLRP1 by inducing FIIND^UPA^-CARD oligomerization.

We have recently demonstrated that primary and immortalized human keratinocytes can undergo rapid NLRP1 inflammasome activation due to high endogenous levels of ASC, caspase-1 and IL-1B (*8, 25*). This provides a robust cellular system to determine if 3Cpros can activate the NLRP1 inflammasome endogenously. Immortalized human keratinocytes were transfected with vector control, HRV14-3Cpro, its catalytically inactive mutant C146A, as well as the constitutively active NLRP1 fragment FIIND^UPA^-CARD, which served as a positive control (Fig. 1E). Like the FIIND^UPA^-CARD fragment, HRV14-3Cpro caused significant IL-1B cleavage and secretion (Fig. 1E,lanes 2 and 3 vs. lane 4). This was accompanied by established hallmarks of NLRP1 activation, including the degradation of the NLRP1 N-terminal fragment and the formation of endogenous ASC oligomers revealed by DSS-crosslinking (Fig. 1E,lanes 2-3, lower panel). Live-cell imaging of human keratinocytes expressing GFP-tagged ASC further corroborated that HRV14-3Cpro expression led to robust ASC-GFP speck formation, which was followed by rapid membrane permeabilization marked by propidium iodide (PI) staining (Fig. 1F). A subset of 3Cpro-transfected cells also underwent apoptosis without ASC speck formation (Fig. 1F,right panel), which was in agreement with the well-established pro-apoptotic roles of 3Cpros (*26, 27*). Taken together, these results demonstrate that 3Cpros can trigger significant inflammasome activation and pyroptosis in immortalized human keratinocytes. CRISPR/Cas9 mediated deletion of *NLRP1, PYCARD*and *CASP1 (25)*(fig. S1C-D) in these cells abrogated the ability for HRV14-3Cpro and HRV16-3Cpro to induce IL-1B secretion (Fig. 1G), proving that 3Cpros trigger a canonical inflammasome response that strictly requires NLRP1, ASC and caspase-1 in human keratinocytes.

Picornaviral 3Cpros are cysteine proteases with well-defined catalytic activity and broad substrate preferences (*20, 28*). Just as anthrax LF cleaves rodent Nlrp1b directly (*14, 15*), we conjectured that 3Cpros could activate human NLRP1 via direct cleavage. Overexpressed NLRP1 undergoes partial auto-cleavage within its FIIND and thus appears as two bands that differ by ∼20 kDa when visualized with an N-terminal fragment specific antibody (*29, 30*) (Fig. 2A,lane 2). In the presence of HRV14-3Cpro, two additional bands became apparent using the same antibody (Fig. 2A,lane 4), while the C-terminal FIIND^UPA^-CARD fragment remained intact, suggesting that the HRV14-3Cpro cleaved NLRP1 close to the N-terminus and not within the FIIND^UPA^-CARD fragment. To visualize the 3Cpro-specific cleavage more accurately, the same experiment was carried out using the NLRP1^F1212A^mutant, which cannot undergo auto-cleavage and thus remains as a single band by SDS-PAGE (Fig. 2B,lane 2). With increasing amounts of HRV14-3Cpro, NLRP1^F1212A^became cleaved into a single proteolytic product, which was approximately 20 kDa smaller than full-length NLRP1 (Fig. 2B). In both experiments, the proteolytic banding patterns could be interpreted by a single cleavage site approximately ∼20 kDa from the NLRP1 N-terminus. We confirmed that the same cleavage could be observed when NLRP1-expressing cell-free lysate was incubated with recombinant HRV14-3Cpro, suggesting that the observed cleavage was most likely direct (Fig. 2C).

**Fig. 2.**
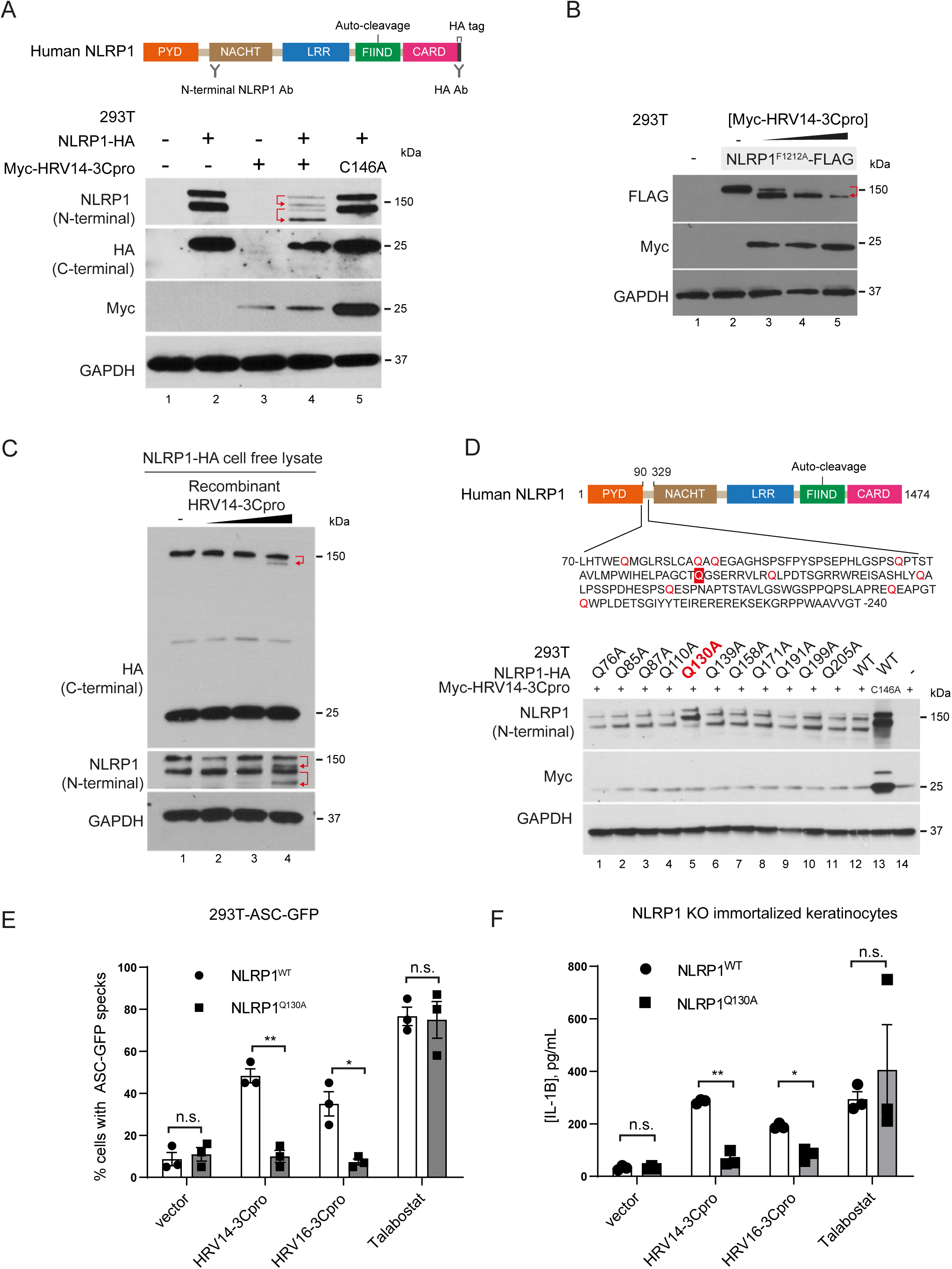
3CPros activate NLRP1 by direct cleavage at a single site. A. HRV14-3Cpro cleaves NLRP1 close to its N-terminus. Top panel: the antibodies used to detect the NLRP1 auto-proteolytic fragments. The epitope of the N-terminal NLRP1 antibody is between NLRP1 a.a. 130 and a.a. 230. Bottom panel: 293T cells were transfected with C-terminally HA-tagged NLRP1 and Myc-tagged HRV14-3Pro. Full length NLRP1 and its cleavage products were B. visualized with the N-terminal fragment-specific NLRP1 antibody and an antibody against the C-terminal HA tag. Red arrows indicate the proposed proteolytic relationship between the observed NLRP1 fragments. Note that the presence of catalytically active 3Cpro decreased the expression all transfected plasmids (see also Fig. 2b, d and fig. S1e). C. HRV14-3Cpro cleaves NLRP1^F1212A^at a single site. 293T cells were transfected with FLAG-tagged NLRP1^F1212A^with increasing amounts of HRV14-3Cpro. Cell lysates were harvested 48 hours post transfection. D. Recombinant HRV14-3Cpro cleaves human NLRP1. Cell-free lysate from NLRP1-HA-transfected 293T cells were incubated with recombinant HRV14-3Cpro at 33°C for 90 mins and analyzed by SDS-PAGE. E. Mapping of the 3Cpro cleavage site. Top: NLRP1 linker region immediately after the PYRIN domain (PYD). Glutamine (Q) residues are highlighted in red. Bottom: 293T cells were co-transfected with the indicated NLRP1 Q>A mutants and HRV14-3Cpro. Total cell lysates were harvested 48 hours post transfection. F. 3Cpros cannot activate the cleavage-resistant NLRP1^Q130A^mutant. 293T-ASC-GFP cells were fixed 24 hours post transfection. The number of cells with ASC-GFP specks were visually scored with wide-field epifluorescence microscopy at 20x maganification. *, p<0.05; **, p<0.005 (Two-way ANOVA). n=3 biological replicates in which more than 100 cells were scored. G. NLRP1^Q130A^cannot restore 3Cpros-dependent IL-1B secretion in *NLRP1*knockout human keratinocytes. *NLRP1*KO keratinocytes were transfected with wild-type NLRP1, NLRP1^Q130A^and 3Cpros. Conditioned were harvested 48 hours post transfection and analyzed by IL-1B ELISA. *, p<0.05; **, p<0.005 (Two-way ANOVA). n=3 biological replicates.

Given the discrete proteolytic products, the 3Cpro cleavage site was mapped to the linker region immediately after the PYRIN domain (PYD) (Fig. 2D,top panel), which is not conserved in rodents. As 3Cpros require a glutamine residue at the P’-1 substrate site (*28*), we mutated all 11 glutamine residues in this linker region to alanine. Of all the mutants tested, only the missense Q130A mutation abrogated NLRP1 cleavage by HRV14-3Cpro (Fig. 2D,lane 5) and other 3CPros (fig. S1E,lanes 1-5 vs lanes 6-10). The Q130A mutation did not affect NLRP1 auto-cleavage within the FIIND (fig. S1E,HA panel). When HRV14-3Cpro and HRV16-3Cpro were co-expressed with NLRP1^Q130A^in 293T-ASC-GFP cells, the 3Cpros could no longer induce ASC-GFP specks formation as they did in wild-type NLRP1 expressing cells (Fig. 2E). Remarkably, NLRP1^Q130A^remained fully sensitive to Talabostat (Fig. 2E), which de-represses DPP8/9-mediated inhibition of the NLRP1-FIIND (*25, 28, 31*). To test the effect of the 3Cpro cleavage site mutation on endogenous NLRP1 inflammasome signaling, wild-type NLRP1 and NLRP1^Q130A^were re-expressed in *NLRP1*knockout immortalized human keratinocytes (fig. S1F,Fig. 2F). While both wild-type NLRP1 and NLRP1^Q130A^restored IL-1B secretion following Talabostat treatment, NLRP1^Q130A^-reconstituted keratinocytes could not secrete IL-1B following transient overexpression of 3Cpro, unlike wild-type NLRP1-reconstituted keratinocytes (Fig. 2F). Therefore, 3Cpros activate the human NLRP1 inflammasome via a different mechanism than Talabostat and relies entirely on a single cleavage site (a.a. Q130-G131) within the human-specific linker region.

To test the effect of live HRV infection on the human NLRP1 inflammasome, NLRP1-HA was stably expressed in HeLa-Ohio cells, which are widely used to study HRV infection. Robust HRV16 viral replication was achieved 16 hours post inoculation (MOI=1) as detected by the accumulation of the viral capsid protein VP2 (Fig. 3A,lanes 4, 5, 10 and 11). This was accompanied by the appearance of the ∼120 kDa NLRP1 cleavage product in infected cells (Fig. 3A,lanes 10 and 11). A small molecule pan-HRV-3Cpro inhibitor, Rupintrivir completely abrogated NLRP1 cleavage (Fig. 3A,lane 12 vs. lanes 10 and 11). Using HeLa-Ohio cells that stably expressed ASC-GFP, we next tested whether HRV16 infection could lead to inflammasome complex assembly. Even though HeLa-Ohio cells support less NLRP1 inflammasome activation than other cell types, HRV16 caused significant ASC-GFP oligomerization (fig. S2A,lane 2 vs. lane 1) and speck formation in cells expressing wild-type NLRP1, but not the cleavage-resistant NLRP1^Q130A^mutant (Fig. 3B,fig. S2A,lane 2 vs. lane 5, fig. S2B). Therefore, in a heterologous system, live HRV16 infection activates the NLRP1 inflammasome and requires direct NLRP1 cleavage.

**Fig. 3.**
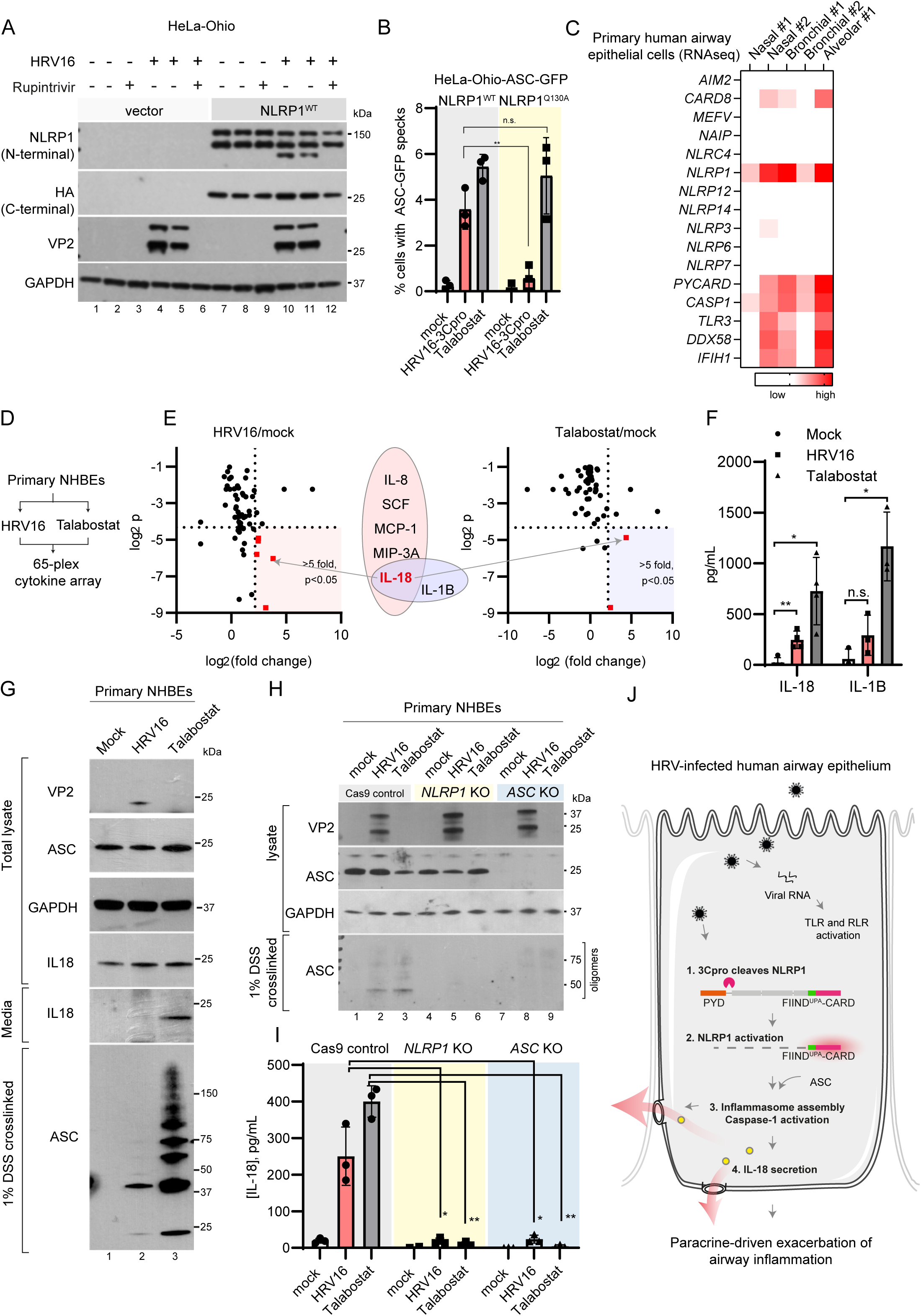
NLRP1 is required for HRV-induced inflammasome assembly and IL-18 secretion in primary human bronchial epithelial cells. A. HRV16 infection causes NLRP1 cleavage. HeLa-Ohio cells stably expressing NLRP1-HA and a vector control were infected with HRV16 at MOI=1 in duplicates. Rupintrivir (10 nM) was added at the time of infection and cell pellets were harvested 24 hours after inoculation. B. The effect of HRV16 infection on ASC-GFP speck formation in HeLa-Ohio-ASC-GFP cells expressing wild-type NLRP1 or NLRP1^Q130A^. ***, p<0.0005 (Two-way ANOVA). n= 3, >100 cells per condition. C. Expression of NLR sensors, inflammasome effectors and RLR sensors in primary human airway epithelial cells. Expression values of Nasal #2 (GSE55458), Bronchial #2 (GSE107971) and Alveolar #2 (GSE83501) and were log-transformed prior to color mapping. D. Comparative analysis of cytokines/chemokines in HRV16-infected and Talabostat-treated NHBEs. Three different lots of NHBEs were used. E. Cytokines/chemokines induced by HRV16 infection and Talabostat in NHBEs. NHBEs were infected with HRV16 (MOI=5) or treated with Talabostat (2 µM). Conditioned media were harvested 48 hours post infection and subjected to multiplex cytokines/chemokine analysis. Cytokines/chemokines that were induced at least 5 fold in each condition (p<0.05, Student’s t test, lognormal values) are highlighted in red. F. HRV16 infection causes robust IL-18 secretion, while IL-1B secretion is variable. Cytokine levels were measured by ELISA. *, p<0.05; **, p<0.005 (Two-way ANOVA). n= 4 biological replicates for IL-18 and 3 biological replicates for IL-1B. G. HRV16 infection causes IL-18 processing and endogenous ASC oligomerization in NHBEs. Cell lysates and conditioned media were harvested 48 hours post HRV16 inoculation. H. HRV16-induced inflammasome assembly requires NLRP1. Cas9-control, *NLRP1*and *ASC*KO NHBEs were infected with HRV16 as in e-g. Cell pellets were harvested 48 hours post infection. I. HRV16-induced IL-18 secretion requires NLRP1 and ASC. Conditioned media were harvested from control and KO NHBEs. IL-18 levels were measured with ELISA. *, p<0.05; **, p<0.005 (Two-way ANOVA), n=3 biological replicates. J. Proposed mechanism of HRV-triggered NLRP1 inflammasome activation and IL-18 secretion in human airway epithelial cells.

HRV is one of the most common causes of human respiratory tract infections and directly infect airway epithelial cells. By examining published RNAseq datasets, we found that primary human airway epithelial cells (*32*–*36*) express robust levels of inflammasome effectors and viral nucleic acid sensors (e.g. *TLR3, RIG-I*and *MDA5*), but a very restricted repertoire of NLR sensors. *NLRP1*was the predominant NLR sensor expressed in all airway epithelial cells examined, whereas other known inflammasome sensors such as *NLRP3*and *AIM2*were expressed at low levels (Fig. 3C,red) or absent. All airway epithelial cell types also express robust levels of endogenous *PYCARD*(ASC), *CASP1*and IL-1 cytokines, *IL1B*and *IL18*, suggesting that they are ‘primed’ to undergo inflammasome activation.

Next we profiled the cytokine and chemokine response of primary human bronchial epithelial cells (NHBEs) to HRV16 infection and Talabostat (Fig. 3D). Remarkably, IL-18, whose secretion strictly depends on inflammasome activation, was the most highly induced cytokine in both HRV16-infected and Talabostat-treated NHBEs (Fig. 3E-F). Although IL-1B secretion was also observed in HRV-infected NHBEs in several experiments, its levels were highly variable between different commercial lots (Fig. 3F). HRV16-infected NHBEs demonstrated other hallmarks of inflammasome activation, such as proteolytic IL-18 processing (Fig. 3G,middle panels), endogenous ASC oligomerization (Fig. 3G,lower panel) and characteristic membrane ‘ballooning’ associated with pyroptosis (fig. S2C). These results demonstrate that HRV16 infection activates the canonical inflammasome pathway in primary bronchial epithelial cells, likely in parallel with other innate immune pathways (Fig. 3E).

Finally, to test whether endogenous NLRP1 is the obligate sensor for HRV-triggered inflammasome activation, we generated *NLRP1*knockout NHBEs. CRISPR/Cas9-mediated deletion of *NLRP1*and inflammasome sensor *ASC*(*PYCARD*) rendered NHBEs resistant to Talabostat-induced pyroptotic cell death marked by propidium iodide staining (fig. S2D). As compared to wild-type NHBEs, HRV16-infected *NLRP1*knockout NHBEs were unable to assemble the inflammasome adaptor ASC into high molecular weight oligomers (Fig. 3H,lanes 2 and 3 vs. 5 and 6). In addition, HRV16-induced IL-18 secretion was abolished in both *NLRP1*and *ASC*(*PYCARD*) knockout NHBEs (Fig. 3I), despite similar levels of viral capsid protein VP2 (Fig. 3H,lane 2, 5 and 8). These results demonstrate that human NLRP1 is an essential, if not the only inflammasome sensor for HRV in primary bronchial epithelial cells. Together with the adaptor protein ASC, NLRP1 is indispensable for HRV-triggered inflammasome activation and subsequent IL-18 secretion.

In summary, we report that enteroviral 3C proteases activate human inflammasome sensor NLRP1 via direct cleavage (Fig. 3J). Together with recent reports, these results provide a unified ‘activation-by-cleavage’ mechanism for the NLRP1 inflammasome in multiple species (*16, 37, 38*). It is noteworthy that even though a single cleavage site mutation (Q130A) can prevent 3Cpros from activating the NLRP1 inflammasome, NLRP1 can still be activated by DPP8/9 inhibitor Talabostat. This suggests that there are at least two independent mechanisms of activation for NLRP1, and the potential of protease-independent NLRP1 activation in exposome defense remains to be fully discovered. Our identification of enteroviral 3Cpros as the ‘missing’ PAMP trigger also sheds light on the evolutionary trajectory of NLRP1. The 3Cpro cleavage site arose in the common ancestor for simian primates (i.e. apes, new world and old world monkeys) and is absent in pro-simians such as tarsiers and lemurs (fig. S3). It is conceivable that the more recently evolved, 3Cpro-responsive *NLRP1*allele has provided a selective survival advantage during the evolution of simian primates including humans.

Our findings establish human NLRP1 as a prominent viral sensor in the airway epithelium, which likely functions in parallel with other immune sensors such as TLRs and RLRs (Fig. 3J). As the inflammasome pathway does not require *de novo*protein synthesis, it may provide a ‘fail-safe’ defense mechanism when host transcription and translation have been shut off by viral virulence factors (*39*). Importantly, our findings demonstrate that the NLRP1 inflammasome is indispensable for IL-18 secretion in HRV-infected primary airway epithelial cells (Fig. 3J). We propose that the NLRP1-IL-18 axis plays a critical role in both antiviral defense and pathological inflammation in the human airway epithelium. IL-18 is known for its ability to stimulate interferon-γ production in T cells, NK cells and MAIT cells (*40, 41*), which is important for the propagation of systemic antiviral response. On the other hand, respiratory viral infections, including HRVs, are well known risk factors for acute exacerbations of asthma and chronic obstructive pulmonary disease (COPD). IL-18 has also been suggested to play a causal role in this context (*42*–*44*). Our findings therefore suggest that the NLRP1 inflammasome-IL-18 axis is a potential therapeutic target to treat inflammatory diseases involving the human airway epithelium.

## Acknowledgements

The authors would like to thank all members of the Reversade laboratory for support and helpful scientific discussion. We also acknowledge the generous support and advice from Dr. Luo Dahai (NTU), Dr. Wu Bin (NTU), Dr. Radoslaw Sobota (IMCB) and Dr. Seth Masters (WEHI). This work is supported by Agency for Science, Technology and Research (GODAFIT Strategic Positioning Fund, Bruno Reversade), National Medical Research Council, Singapore (NMRC/OFYIRG/0046/2017, Franklin Zhong), Concern Foundation’s Conquer Cancer Now Award (Franklin Zhong) and National Research Foundation, Singapore (Franklin Zhong). Bruno Reversade is a fellow of the Branco Weiss Foundation and a recipient of the A*STAR Investigatorship, the senior NRF investigatorship and EMBO Young Investigatorship. Franklin Zhong is a Nanyang Assistant Professor and National Research Foundation Fellow.

## Author contributions

F.L.Z., K.S.R. and D.E.T.T. initiated, designed, performed and analyzed the experiments. T.K.S., O.H.H. and L.T.S. performed all viral infection experiments. C. L. performed the Luminex assay. V.T.K.C. and W.D.Y. supervised virology experiments. F.L.Z. and B.R. jointly conceived of the study, supervised experiments and data analysis and wrote the manuscript.

## Supplementary Materials

### Methods

#### Cell culture

293Ts (ATCC #CRL-3216), HeLa-Ohio (ECACC General Cell Collection #84121901) and normal bronchial epithelial cells (Lonza #CC-2541) were obtained from commercial sources and cultured according the suppliers’ protocols. Immortalized human keratinocytes (N/TERT-1) were a kind gift from H. Reinwald (MTA) (*45*). All cell lines underwent routine Mycoplasma testing with Lonza MycoAlert (Lonza #LT07-118).

#### Plasmid transfection and stable cell line generation using lentiviruses

293T-ASC-GFP and N/TERT-ASC-GFP were described previously (*25*). All transient expression plasmids were cloned into the pCS2+ vector using standard restriction cloning. Polyclonal Cas9/CRISR knockout cell lines were generated with lentiCRISPR-v2 (Addgene #52961) and selected with puromycin. Knockout efficiency was tested with Western blot 7-10 days after puromycin selection. Site-directed mutagenesis was carried out with QuickChangeXL II (Agilent #200522).

#### Antibodies and cytokine analysis

The following antibodies were used in this study: c-Myc (Santa Cruz Biotechnology #sc-40), HA tag (Santa Cruz Biotechnology, #sc-805), GAPDH (Santa Cruz Biotechnology, #sc-47724), ASC (Apipogen, #AL-177), CASP1 (Santa Cruz Biotechnology, #sc-622), IL1B (R&D systems, #AF-201), FLAG (SigmaAldrich, #F3165), GFP (Abcam, #ab290), NLRP1 (R&D systems, #AF6788), IL18 (R&D systems, MAB9124) and VP2 (QED Bioscience, #18758). Cytokine and chemokine measurements were carried out with human IL-1B ELISA kit (BD, #557953), human IL-18 ELISA kit (MBL Bioscience, #7620) and Immune Monitoring 65-Plex Human ProcartaPlex™ Panel (ThermoFisher EPX650-10065-901).

### HRV16 virus propagation

HRV used in the study was HRV-A16 (strain 11757; ATCC VR-283, Manassas, VA, USA), and was propagated in HeLa cell line (HeLa Ohio, ECACC 84121901, Porton Down, Salisbury, Wiltshire, UK). HeLa cells were grown in Eagle’s Minimum Essential Medium (EMEM) ATCC® 30-2003™, supplemented with 10% fetal bovine serum (FBS) (BioWest, Kansas City, MO, USA), 2% HEPES and 1% Antibiotic-Antimycotic (Anti-Anti) (Gibco) and incubated at 37°C humidified incubator with 5% CO2. To propagate HRV16, HeLa cells were first seeded to achieve confluency of about 80-90% in 24-well plate overnight (Figure 2A). Cells were rinsed with 1X dulbecco’s phosphate-buffered saline (dPBS) and infected with HRV16 before addition of EMEM with 2% FBS, 2% HEPES and 1% Anti-Anti. Infected HeLa cells were incubated at 33°C for two to three days. Viruses were harvested from the supernatants of infected HeLa cells when about 80% cytopathic effects (CPE) was observed (Figure 2B). HRV virus stocks were centrifuged at 3500rpm for 10mins at 4°C to remove cellular debris, and aliquoted into cryovials for storage at −80°C.

### Inoculation of human rhinovirus

HRV was diluted using the respective cell culture medium and inoculated at multiplicity of infection (MOI) of 5.0 (NHBE) and 1.0 (HeLa), respectively. Infected cells were incubated at 33 °C for 1 hr. Conditioned non-infected cell culture medium from viral propagation was added as uninfected-control. The HRV-infected and control cells were then incubated at 33 °C for up to 96 hours post-infection (hpi). Cell culture supernatant and cell lysate were collected to perform relevant assays between 24 − 72 hpi.

### Viral quantification using rhinovirus plaque assay

HeLa cells (at 85-95% confluence) in 24-well plates were incubated with 100 µL of serial dilutions from 10^−1^to 10^−6^of sample from infected hNECs at 33 °C for 1 h. The plates were rocked every 15 min to ensure equal distribution of virus. The inoculum was removed and replaced with 1 mL of Avicel (FMC Biopolymer) overlay to each well, and incubated at 33 °C for 65-72 h. The overlay components were optimized to obtain HRV plaques suitable for counting. Avicel powder was added into double strength MEM to formulate 1.2% Avicel solution, and with a final concentration comprising 3% FBS, 2% HEPES, 1.5% NaHCO3, 3% MgCl2 and 1% Anti-Anti. Avicel overlay was removed after the incubation period, and cells were fixed with 20% formalin in 1× PBS for 1 h. Formalin was removed, and cells were washed with 1× PBS. The fixed cells were stained with 1% crystal violet for 15 min, and washed. The plaque-forming units (PFU) were calculated as follows: Number of plaques × dilution factor = number of PFU per 100 µL, which is then expressed as PFU/mL.

## Legends for Fig. S1-S3

**Fig. S1.**
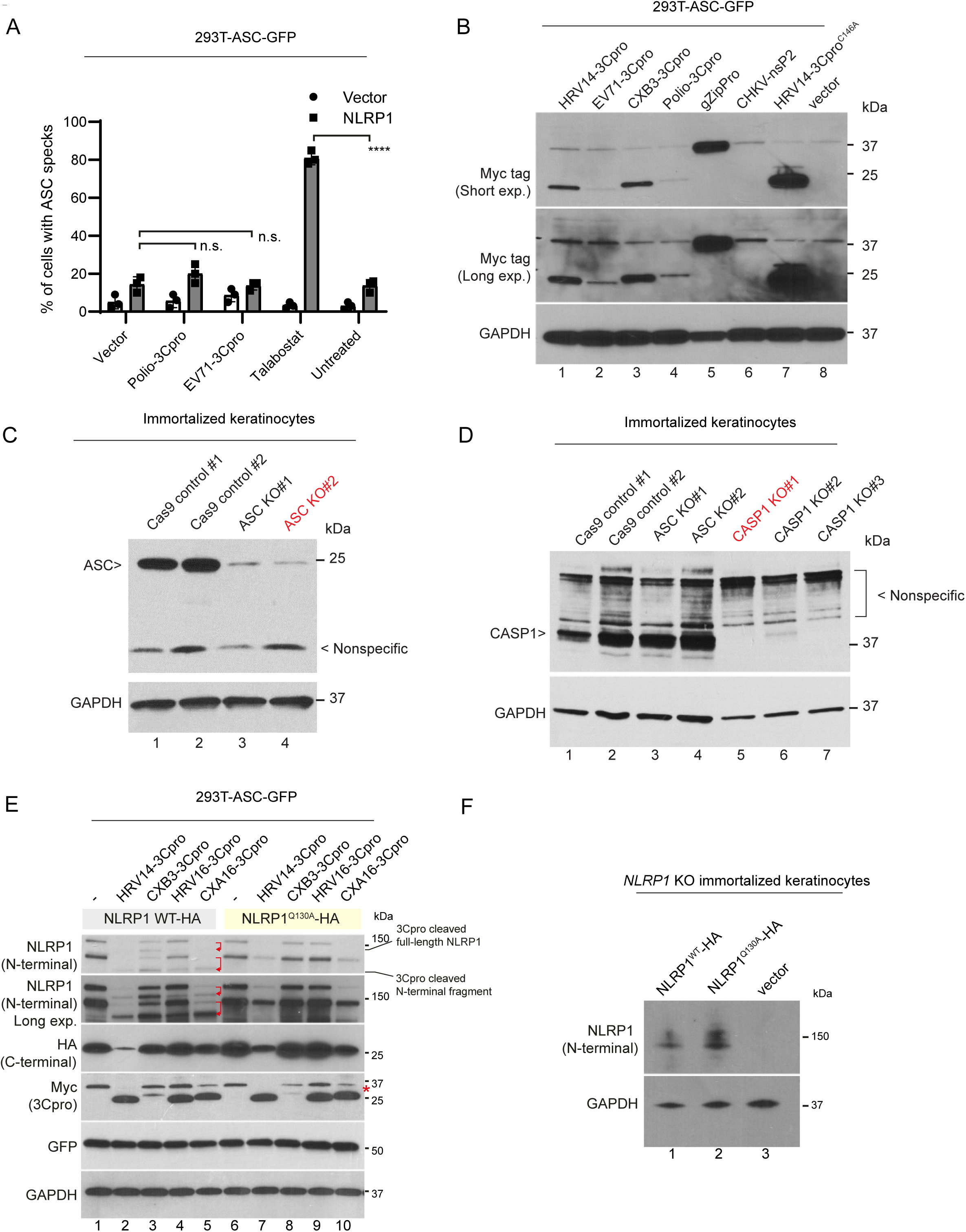
A. Polio- and EV71-3Cpros do not significantly induce ASC-GFP speck formation in 293T-ASC-GFP cells expressing NLRP1. B. Expression levels of viral proteases in 293T-ASC-GFP cells. All indicated viral proteases were tagged with Myc at the N-terminus. Cell pellets were harvested 48 hours post plasmid transfection. C. Validation of pooled CRISPR/Cas9 *PYCARD/ASC*knockout in immortalized keratinocytes. The cell lines that were used for downstream analyses are highlighted in red. D. Validation of pooled CRISPR/Cas9 *CASP1*knockout in immortalized keratinocytes. E. NLRP1^Q130A^is resistant to cleavage by multiple 3Cpros. 293T-ASC-GFP cells were transfected as in Fig. 2d. Cell pellets were harvested 48 hours post transfection. Asterisk, endogenous c-myc in 293T cells (unrelated to myc-3Cpros). F. NLRP1^WT^-HA and NLRP1^Q130A^-HA were expressed to similar levels in *NLRP1*KO immortalized keratinocytes.

**Fig. S2.**
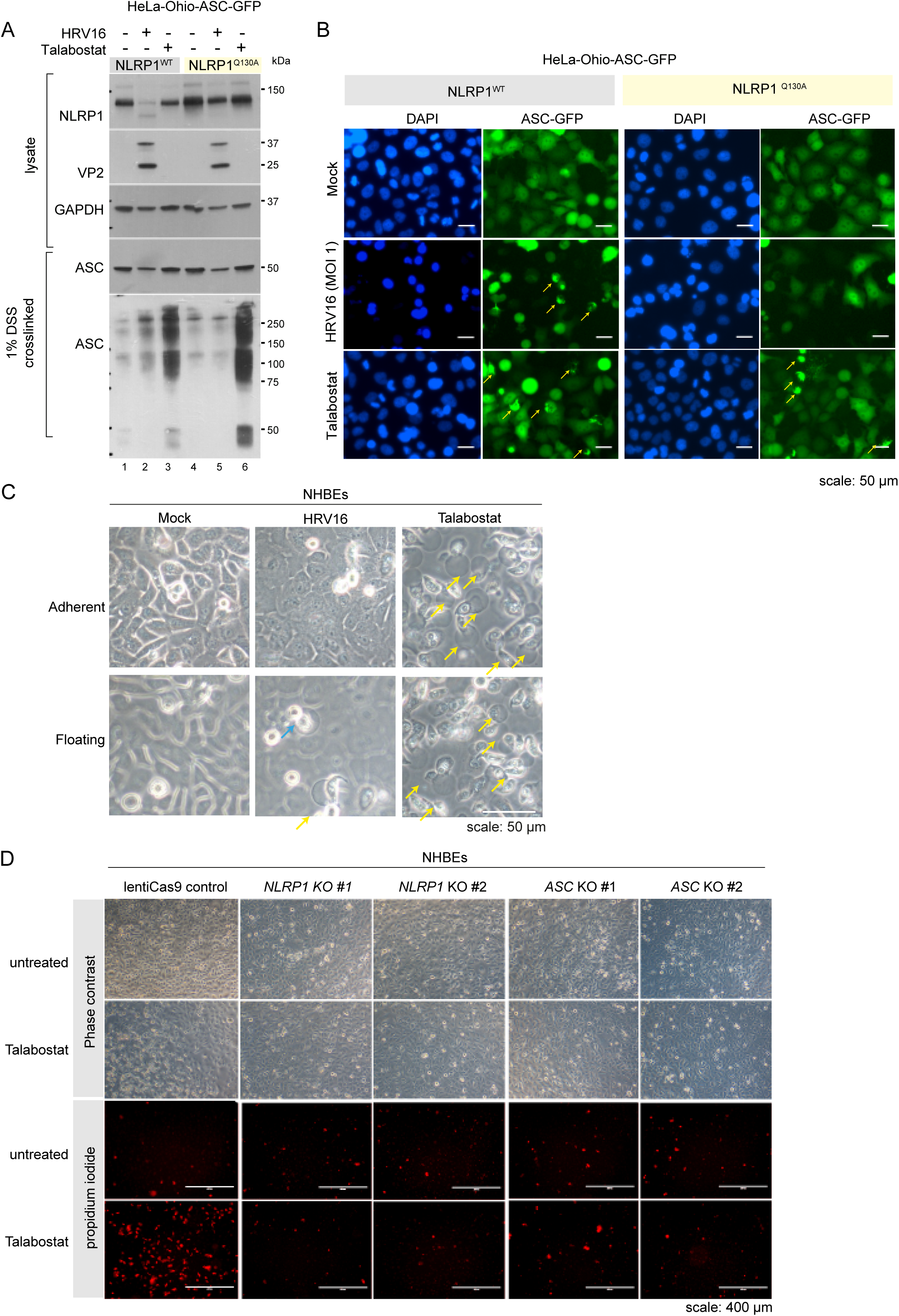
A. HRV16 causes ASC-GFP oligomerization in a cleavage-dependent manner. HeLa-Ohio-ASC-GFP reporter cells were stably transduced with wild-type NLRP1 or the cleavage-resistant NLRP1^Q130A^mutant and infected with HRV16 (MOI=1) as in a. ASC-GFP oligomers were crosslinked with 1% DSS and solubilized with 1% SDS. Note that level of overall ASC-GFP was reduced in HRV16-infected cells. B. The effect of HRV16 infection and Talabostat on HeLa-Ohio-ASC-GFP cells expressing NLRP1. Yellow arrows, ASC-GFP specks. C. Morphology of cell death in HRV16-infected NHBEs. Yellow arrows, characteristic pyroptotic cell death (thistle-like, elongated cell bodies with membrane ‘balloons’). Blue arrows, late-stage apoptosis (membrane blebs with shrunken cell bodies). D. Functional validation of CRISPR/Cas9 knockout NHBEs. Cell death was visualized by propidium iodide staining.

**Fig. S3.**
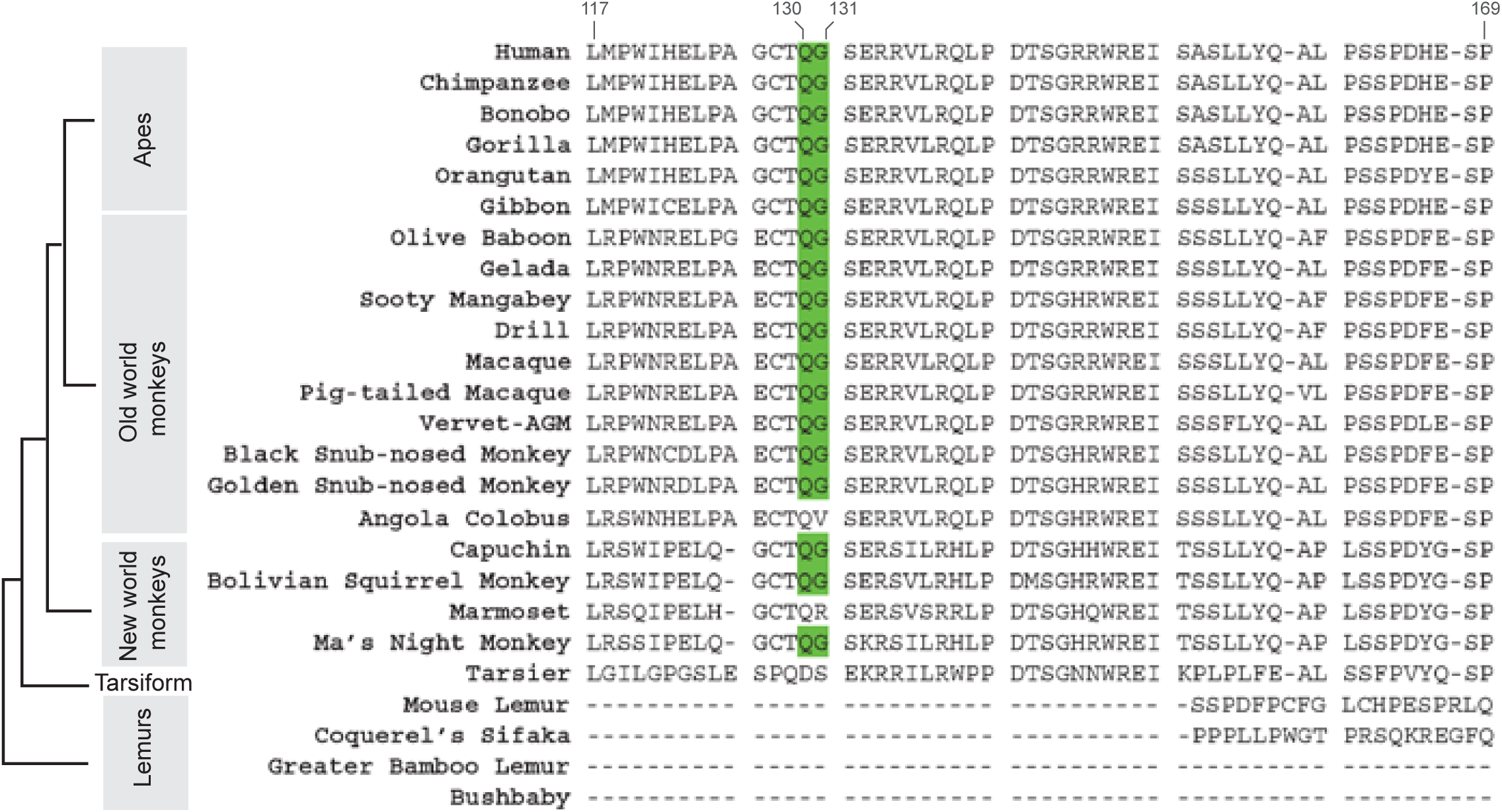
Multiple sequence alignment of the linker region in primate NLRP1 homologs. The 3Cpro cleavage site (p. Q130-G131 in human NLRP1) is highlighted in green.

